# Endothelial cell senescence exacerbates pulmonary hypertension through Notch-mediated juxtacrine signaling

**DOI:** 10.1101/2021.02.02.429321

**Authors:** Risa Ramadhiani, Koji Ikeda, Kazuya Miyagawa, Gusty Rizky Teguh Ryanto, Naoki Tamada, Yoko Suzuki, Ken-ichi Hirata, Noriaki Emoto

## Abstract

Pulmonary arterial hypertension (PAH) is a fatal disease characterized by pathological pulmonary artery remodeling. Endothelial cells (EC) injury including DNA damage is critically involved in the vascular remodeling in PAH, and persistent injury leads to cellular senescence in ECs. Here, we show that EC senescence exacerbates pulmonary hypertension through Notch-mediated juxtacrine signaling. EC-specific progeroid mice that we recently generated showed exacerbated pulmonary hypertension after chronic hypoxia exposure, accompanied by the enhanced pulmonary arterial smooth muscle cells (PASMCs) proliferation in the distal pulmonary arteries. Mechanistically, we identified that senescent ECs highly expressed Notch ligands, and thus activated Notch signaling in PASMCs, leading to enhanced PASMCs proliferation and migration capacities. Consistently, pharmacological inhibition of Notch signaling attenuated the effects of senescent ECs on SMCs functions *in vitro*, and on the pulmonary hypertension in EC-specific progeroid mice *in vivo*. These data establish EC senescence as a crucial disease-modifying facor in PAH.

## Introduction

Pulmonary arterial hypertension (PAH) is a devastating lung disease characterized by progressive vasculopathy of small pulmonary arteries, leading to elevated pulmonary arterial pressure and right heart failure (Rabinovitch, 2012; Thompson and Lawrie, 2017). The recent development of new drugs for PAH significantly improved patients’ quality of life, hemodynamic parameters, and clinical outcomes. However, long-term prognosis remains unsatisfactory with 5-years survival of ~65% (Vachiéry and Gaine, 2012; Farber *et al.*, 2015; Grinnan *et al.*, 2019; Prins *et al.*, 2019). Post mortem analysis of the lungs from PAH patients whose symptoms were well-controlled on prostacyclin analog, showed extensive plexogenic arteriopathy, a hallmark of PAH (Rich *et al.*, 2010). These findings indicate that disease progression is inevitable throughout currently available therapies, resulting in poor long-term survival in PAH. Thereby, the development of new therapies targeting the pathological vascular remodeling to improve the vasculopathy is urgently needed.

Several animal studies revealed that hemodynamic unloading reverses the pathological occlusive lesion in pulmonary hypertension (O’Blenes *et al.*, 2001; Mercier *et al.*, 2009; Abe *et al.*, 2016). These studies are consistent with the clinical findings of the regression of vasculopathy lesion after unilateral lung transplantation in the non-transplanted lung (Levy *et al.*, 2001; Deb *et al.*, 2006). In the case of PAH associated with congenital heart disease (PAH-CHD), hemodynamic unloading through shunt closure restores the pulmonary artery pressure and reverses the occlusive arteriopathy lesions in a limited time. However, hemodynamic correction failed to maintain the lesion regression, and eventually irreversible phenotypes that resemble neointimal and plexiform lesions of PAH occur after a certain period. Cellular senescence has recently been revealed as the culprit of reversibility loss in PAH associated with hemodynamic abnormality (Hsu *et al.*, 2016; van der Feen *et al.*, 2017; Van Der Feen *et al.*, 2020). Moreover, the potential link of vascular cell senescence in the pathophysiology of PAH has been recently explained (Van Der Feen, Berger and Bartelds, 2019).

Cellular senescence was initially defined as a stable cell cycle arrest as a consequence of the limited proliferation capacity of cells, i.e. replicative senescence. The emerging evidence suggests another type of senescence, i.e. premature senescence that is a stress response triggered by various stimuli (Herranz and Gil, 2018). Both type of senescent cells have been considered as a driver of age-related disease due to their ability to alter tissue homeostasis and promote secondary senescence through the senescence-associated secretory phenotype (SASP) (Childs *et al.*, 2015; Lecot *et al.*, 2016; Mitri and Alimonti, 2016). Recent studies showed that SASP is not the sole mediator of non-autonomous functionality of cellular senescence, and Notch signaling is also involved in the secondary senescence (Parry *et al.*, 2018; Teo *et al.*, 2019). Here, we explored the potential role of EC senescence in the pathogenesis of PAH and identified a detrimental effect of senescent EC on the progression of pulmonary hypertension through the activation of Notch-mediated juxtacrine signaling in pulmonary smooth muscle cells.

## Results

### EC-specific progeroid mice exhibit the exacerbated pulmonary hypertension

We recently generated EC-specific progeroid mice that overexpress the dominant-negative form of telomeric repeat-binding factor 2 (TRF2DN) under the control of Tie2 or vascular endothelial cadherin (VEcad) promoter (Barinda *et al.*, 2020). By utilizing these EC-specific progeroid mice, we investigated the role of EC senescence in pulmonary hypertension (PH). Under the normoxic condition, there were no differences in pulmonary arterial pressure and systemic hemodynamics between WT and VEcad-TRF2DN-Tg mice (Sup. Fig. 1A–C). After three weeks of exposure to hypoxia (10% O_2_), VEcad-TRF2DN-Tg mice exhibited worsened PH, indicated by the higher right ventricular systolic pressure (RVSP) and augmented right ventricular mass normalized by left ventricle + septum mass (Fulton’s index) (Fig. 1A and 1B). Histological analysis of the lungs showed further reduction in distal pulmonary arteries (PAs) (Fig. 1C and 1D) and deteriorated medial thickening in small PAs (Fig. 1E and 1F) in VEcad-TRF2DN-Tg mice comparing to those in WT mice after chronic hypoxia exposure . Of note, the number of medial proliferating smooth muscle cells (SMCs) in small PAs was significantly increased in the lungs of VEcad-TRF2DN-Tg mice as compared to that in WT mice (Fig. 1G and 1H). These data suggest that EC senescence exacerbates PH potentially through the enhanced SMCs proliferation.

**Figure 1.**
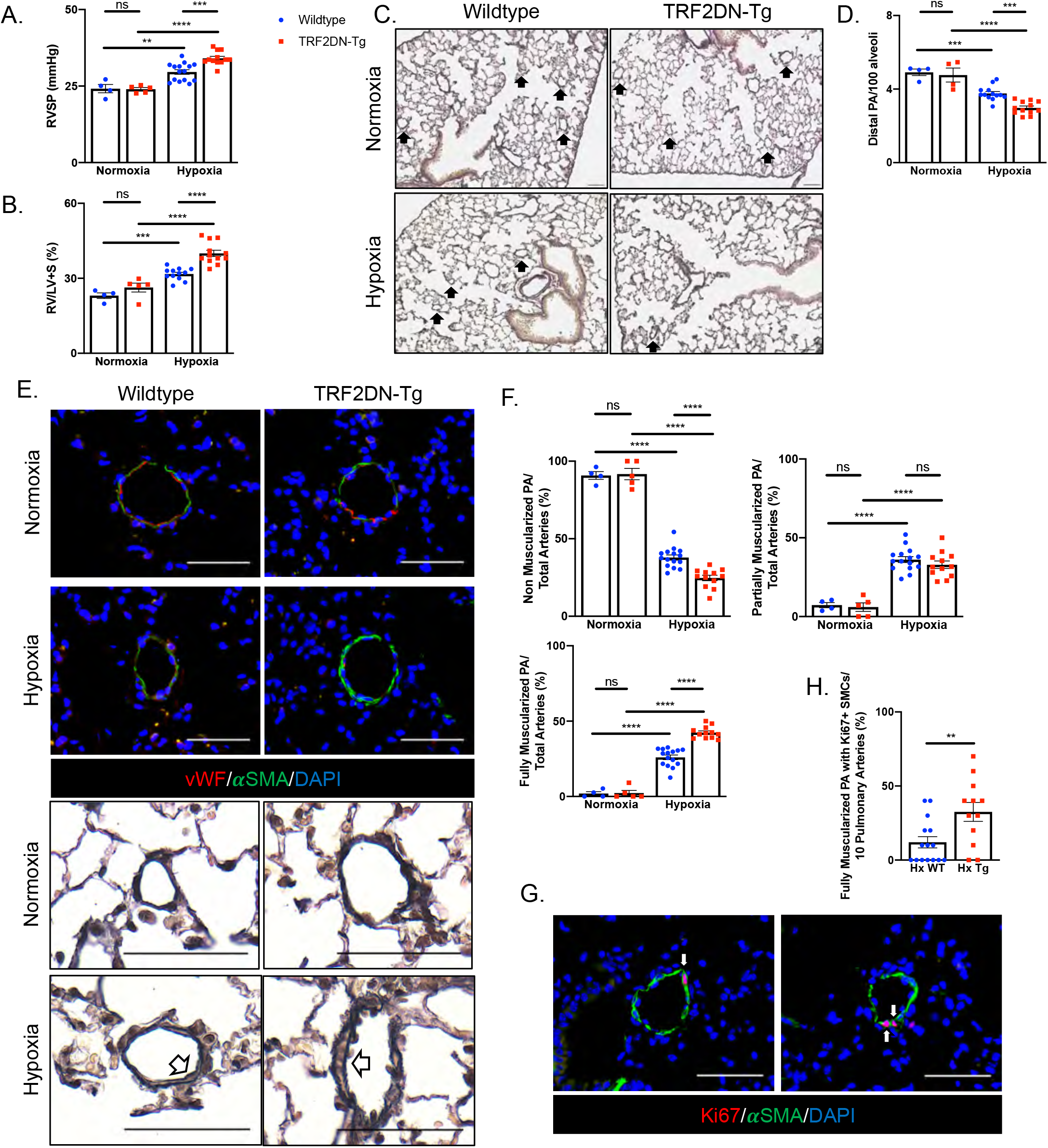
EC-specific progeria exacerbates pulmonary hypertension in mice. (**A**, **B**) Right ventricular systolic pressure (RVSP) (A) and Ratio of right ventricle compared to left ventricle + septum (B) in WT and TRF2DN-Tg mice exposed to either normoxia or hypoxia for 3 weeks. (**C**) Representative images of the lung sections stained with Elastica van gieson. Distal pulmonary arteries (PAs) are indicated by arrowheads. The Lungs dissected from WT and Tg mice exposed to either normoxia or hypoxia were analyzed. (**D**) Quantitative analysis for the number of PAs in the lungs of WT and Tg mice exposed to either normoxia or hypoxia. (**E**) Immunohistochemistry for von Willebrand factor (vWF) and α-smooth muscle actin (αSMA) in the lungs of WT and Tg mice exposed to either normoxia or hypoxia (upper images). Images of the lung sectioned stained with Elastica van gieson were also shown (lower images). Distal PAs with medial thickening were indicated by arrows. (**F**) Quantification of non-, partial-, and fully-muscularized distal PAs in the lungs of WT and Tg mice exposed to either normoxia or hypoxia. (**G**) Immunohistochemistry for Ki-67 and αSMA in the lungs of WT and Tg mice exposed to hypoxia. Arrows indicate the Ki-67-positive proliferating smooth muscle cells (SMCs). (**H**) Quantitative analysis for Ki-67-positive SMCs among 10 fully-muscularized PAs. Data are presented as mean *±* SEM. Two-tailed student’s *t*-test was used for the analysis of the differences between two groups. Two-way ANOVA with Tukey’s post hoc test was used for the analysis of the differences between groups more than three. The number of samples was: n = 4 for normoxic WT; n = 4-5 for normoxic Tg; n = 12-15 for hypoxic WT; n = 12 for hypoxic Tg. Scale bars: 50 μM. *\*P < 0.05, **P < 0.01, ***P < 0.001, and ****P < 0.0001.*

### Senescent ECs promote SMCs proliferation and migration via cell-cell contact

To explore the molecular mechanisms underlying the exacerbated PH in EC-specific progeroid mice, we prepared senescent ECs by overexpressing the TRF2DN *in vitro*. Cellular senescence was validated by increased CDK inhibitors and SASP factors expressions as compared to those in GFP-transfected control cells (Fig. 2A). As medial thickening with increased proliferation of SMCs was prominent in VEcad-TRF2DN-Tg mice exposed to hypoxia, we investigated the effect of senescent ECs on SMCs functions. ECs and SMCs were co-cultured in two conditions; direct (in the presence of cell-cell contact) and indirect (in the absence of cell-cell contact) (Fig. 2B). It has been reported that direct interaction with ECs promotes the contractile phenotype of SMCs (Brown *et al.*, 2005), and we confirmed morphological changes into a spindle shape and increased differentiation markers in pulmonary artery (PA) SMCs directly co-cultured with ECs (Sup. Fig. 2A and 2B). Of note, PASMCs directly co-cultured with senescent PAECs showed enhanced proliferation and migration capacities comparing to PASMCs directly co-cultured with control GFP-transfected ECs (Fig 2C and 2D). Notably, these effects were not found in the indirect co-culture condition (Fig. 2C and 2D). PASMCs apoptosis was not significantly affected by the co-culture with senescent ECs both in direct and indirect conditions (Fig. 2E). These data strongly suggest that the cell-cell contact-mediated interaction between senescent ECs and SMCs enhances the proliferation and migration capacities in PASMCs.

**Figure 2.**
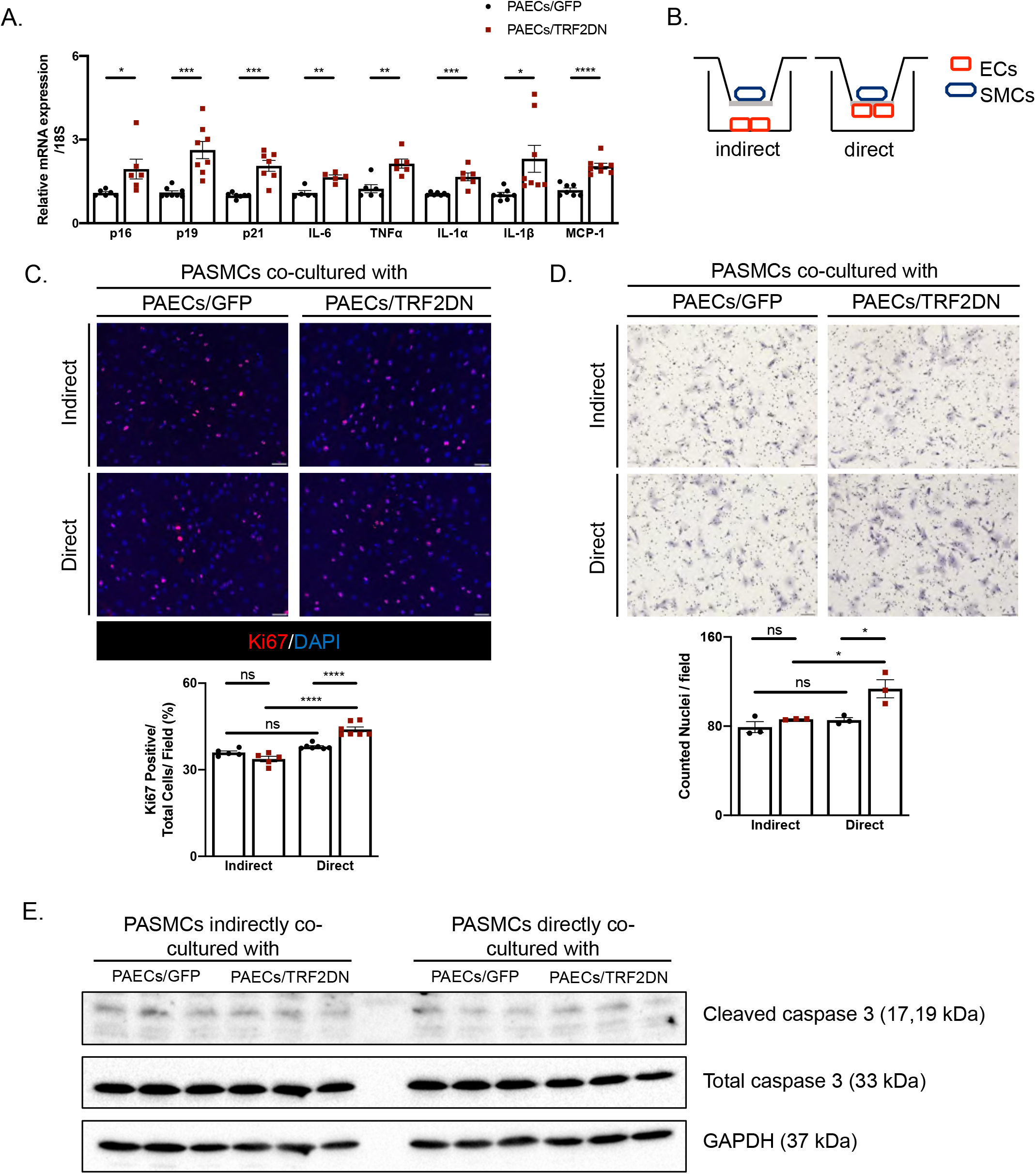
Senescent ECs enhance SMCs proliferation and migration through direct cell-cell contact. (**A**) Real-time qPCR analysis for CDKIs and SASP factors in pulmonary artery ECs (PAECs) transfected with either GFP (control cells) or TRF2DN (premature senescent cells) (n = 5-8 each). (**B**) Schemes for co-culture experiments of PAECs and pulmonary artery SMCs (PASMCs). (**C**) Immunocytochemistry for Ki-67 in PASMCs directly (n = 7 each) or indirectly (n = 5 each) co-cultured with control or premature senescent PAECs. (**D**) Migration capacity was assessed by a modified Boyden chamber assay in PASMCs directly or indirectly co-cultured with control or premature senescent PAECs (n = 3 each). (**E**) Immunoblotting for cleaved caspase-3, total caspase-3, and GAPDH in PASMCs directly or indirectly co-culture PASMCs with control or premature senescent PAECs. Apoptosis was induced by incubating with 500 nM hydrogen peroxide for 3 h (n = 3 each). Data are presented as mean *±* SEM. Two-tailed student’s *t*-test was used for the analysis of the differences between two groups. Two-way ANOVA with Tukey’s post hoc test was used for the analysis of the differences between groups more than three. Scale bars: 50 μM. *\*P < 0.05, **P < 0.01, ***P < 0.001, and ****P < 0.0001.*

### Senescent PAECs enhances PASMCs migration and proliferation through Notch-mediated juxtacrine signaling

Senescent cells have unique non-cell-autonomous functions through senescence-associated secretory phenotype (SASP) that acts in an autocrine or paracrine fashion (Gonzalez-Meljem *et al.*, 2018; Shakeri *et al.*, 2018). Recently, Notch signaling has been identified as a mediator of senescent cells in a cell-contact-dependent juxtacrine fashion (Parry *et al.*, 2018; Teo *et al.*, 2019). We, therefore, investigated the Notch signaling in the communication between senescent ECs and SMCs. Expression of Notch ligands such as Jagged-1, Jagged-2, and Delta-4 were significantly increased in senescent ECs compared to control ECs (Fig. 3A). It has been known that global patterns of DNA hypomethylation are observed in senescence cells, which is associated with the altered expression of many genes (Cruickshanks *et al.*, 2013). Pharmacological inhibition of DNA methylation using 5-azacytidine (5-AZA) partially abolished the increased Notch ligands expression in senescent ECs (Sup. Fig. 2C). These data suggest a potential role of senescence-associated epigenetic modifications in the dysregulated Notch ligands expressions in senescent ECs.

**Figure 3.**
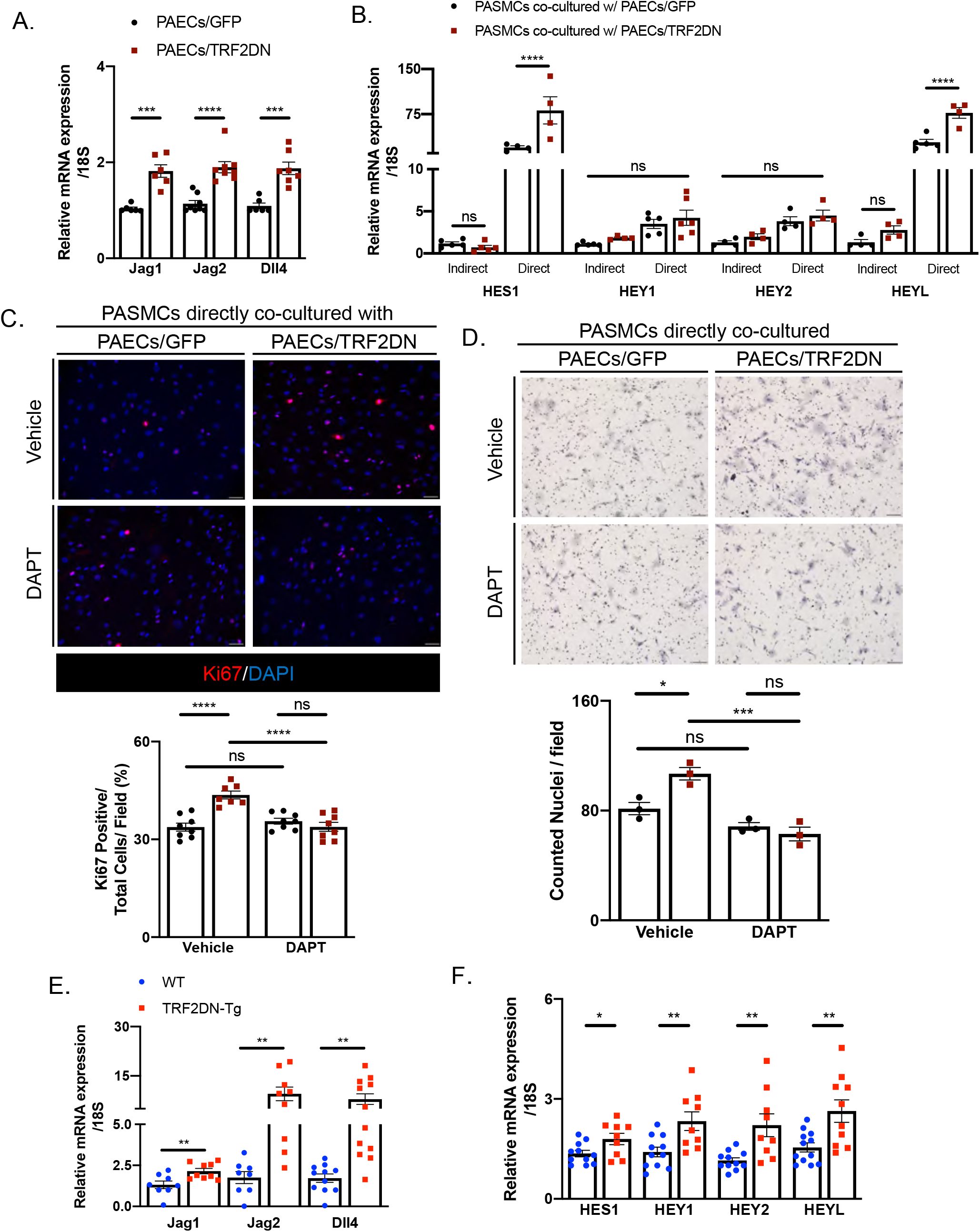
Senescent ECs enhance SMCs proliferation and migration through Notch-mediated juxtacrine signaling. (**A**) Real-time qPCR analysis for Notch ligands in PAECs transfected with either GFP (control cells) or TRF2DN (premature senescent cells) (n= 6-8 each). (**B**) Real-time qPCR for Notch target genes in PASMCs directly or indirectly co-cultured with control or premature senescent PAECs (n = 4-5 each). (**C**) Immunocytochemistry for Ki-67 in PASMCs directly co-cultured with control or premature senescent PAECs. Cells were treated with either vehicle or 10 μM DAPT (n = 7-8 each). (**D**) Migration capacity was assessed by a modified Boyden chamber assay in PASMCs directly co-cultured with control or premature senescent PAECs. Cells were treated with either vehicle or 10 μM DAPT (n= 3 each). (**E**) Real-time qPCR analysis for Notch ligands in ECs isolated from the lungs of WT (n = 8-11) and TRF2DN-Tg (n = 9-10) mice exposed to chronic hypoxia. (**F**) Real-time qPCR analysis for Notch target genes in the lungs of WT (n = 11-12) and TRF2DN-Tg (n = 9-10) mice exposed to chronic hypoxia. Data are presented as mean *±* SEM. Two-tailed student’s *t*-test was used for the analysis of the differences between two groups. Two-way ANOVA with Tukey’s post hoc test was used for the analysis of the differences between groups more than three. Scale bars: 50 μM. *\*P < 0.05, **P < 0.01, ***P < 0.001, and ****P < 0.0001*.

Consistently with the increased Notch ligands expression, more induction of Notch target genes including HES1 and HEYL was detected in PASMCs directly co-cultured with senescent ECs than in SMCs directly co-cultured with control ECs (Fig. 3B). The γ-secretase inhibitor, DAPT, abrogated the enhanced proliferation and migration in PASMCs directly co-cultured with senescent PAECs (Fig. 3C and 3D). These data further support the critical role of Notch-mediated juxtacrine signaling in the augmented capacities of proliferation and migration in PASMCs directly co-cultured with senescent PAECs.

### Inhibition of Notch signaling improves the PH phenotypse in EC-specific progeroid mice

In a way similar to the *in vitro* findings, increased Notch ligands expression, such as Jagged-1, Jagged-2, and Delta-4, were detected in pulmonary ECs isolated from VEcad-TRF2DN-Tg mice. Consistently, the expression of Notch target genes such as HES1, HEY1, HEY2, and HEYL, in the lungs of VEcad-TRF2DN-Tg mice were significantly higher than that in WT mice (Fig. 3E and 3F).

To elucidate an important role of Notch signaling in the exacerbated PH phenotype in EC-specific progeroid mice, we subcutaneously injected either vehicle or DAPT three times a week during chronic hypoxia exposure, as shown in the scheme (Fig. 4A). Beforehand, pharmacological inhibition of Notch signaling was confirmed by the reduced Notch target genes expression in the lungs of WT mice treated with DAPT using this experimental protocol (Fig. 4B). DAPT-treatment ameliorated the PH phenotypes in both WT and VEcad-TRF2DN-Tg mice, signified by lower RVSP and decreased Fulton’s index ratio (Fig. 4C and 4D). Of note, the deteriorated PH phenotypes in VEcad-TRF2DN-Tg mice disappeared after DAPT-treatment (Fig. 4C and 4D). Histological analysis of the lungs also revealed that DAPT-treatment alleviated the worsened pulmonary artery remodelings in VEcad-TRF2DN-Tg mice (Fig. 4E–4H). We confirmed that Notch target genes expression in the lungs was similar between WT and VEcad-TRF2DN-Tg after treatment with DAPT (Fig 4I). These data collectively indicate that senescent ECs deteriorate PH by enhancing SMCs proliferation and migration capacities via Notch-mediated juxtacrine signaling (Fig. 4J).

**Figure 4.**
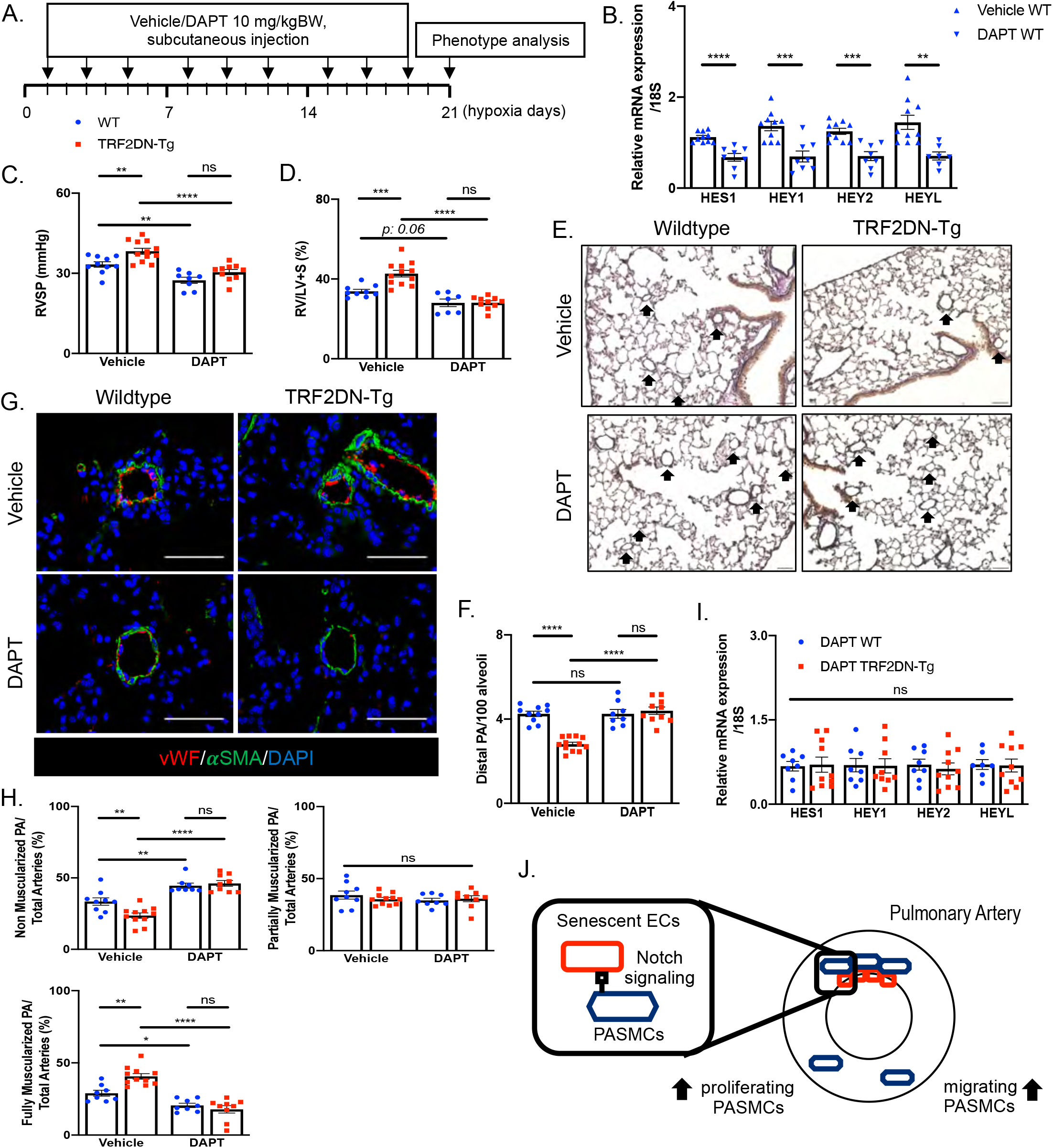
Inhibition of Notch signaling abolishes the exacerbated PH phenotype in EC-specific progeroid mice. (**A**) Schematic diagram for mice experiments. (**B**) Real-time qPCR analysis for Notch target genes in whole lungs of WT mice treated with either vehicle or DAPT according to the experimental protocol shown in A. (**C**, **D**) RVSP (C) and Ratio of right ventricle compared to left ventricle + septum (D) in hypoxia-exposed WT and TRF2DN-Tg mice treated with either vehicle or DAPT. (**E**) Representative images of the lung sections stained with Elastica van gieson. The lungs dissected from hypoxia-exposed WT and TRF2DN-Tg mice treated with either vehicle or DAPT were analyzed. Arrows indicate distal PAs (**F**) Quantitative analysis for the number of distal PAs in the lungs of hypoxia-exposed WT and TRF2DN-Tg mice treated with either vehicle or DAPT. (**G**) Immunohistochemistry for vWF and αSMA in the lungs of WT and TRF2DN-Tg mice treated with either vehicle or DAPT. (**H**) Quantitation for non-, partially, or fully-muscularized distal PAs in the lungs of hypoxia-exposed WT and TRF2DN-Tg mice treated with either vehicle or DAPT. (**I**) Real-time qPCR analysis for Notch target genes in whole lungs of hypoxia-exposed WT and TRF2DN-Tg mice treated with DAPT. (**J**) Schematic diagram for the mechanism underlying the detrimental role of senescent ECs in PH. Data are presented as mean *±* SEM. Two-tailed student’s *t*-test was used for the analysis of the differences between two groups. Two-way ANOVA with Tukey’s post hoc test was used for the analysis of the differences between groups more than three. The number of samples was: n = 9-10 for vehicle-treated WT; n = 11-12 for vehicle-treated Tg; n = 7-8 for DAPT-treated WT; n = 9-10 for DAPT-treated Tg. Scale bars: 50 μM. *\*P < 0.05, **P < 0.01, ***P < 0.001, and ****P < 0.0001.*

## Discussion

In this study, we have revealed a crucial role of senescent ECs in the progression of PAH through the interaction with PASMCs via Notch-mediated juxtacrine signaling. EC senescence has been considered to play a causative role in age-related metabolic (Shosha *et al.*, 2018; Barinda *et al.*, 2020) and cardiovascular disease (Minamino *et al.*, 2002; Ahmad *et al.*, 2015; Yamazaki *et al.*, 2016; Shakeri *et al.*, 2018; Song *et al.*, 2020). Moreover, the potential role of cellular senescence has been explored in various non-age-related diseases such as type-1 diabetes (Thompson *et al.*, 2019) and PAH associated with congenital heart disease (Van Der Feen *et al.*, 2020). Senescence-associated secretory phenotype (SASP) has been highlighted as a non-cell-autonomous activity to bring detrimental effects to nearby cells (Tchkonia *et al.*, 2013; Van Der Feen, Berger and Bartelds, 2019; Li and Lerman, 2020). Recently, Notch-mediated signaling has also been revealed as non-cell-autonomous functionality of cellular senescence in addition to the SASP (Hoare *et al.*, 2016; Parry *et al.*, 2018; Teo *et al.*, 2019).

Senescent cells exhibit a massive transcriptional dysregulation as a consequence of epigenetic modifications including the global DNA hypomethylation, which may promote specific transcriptional programs (Hernando-Herraez *et al.*, 2019). Previous studies revealed that various stimuli transiently or persistently activates Notch signaling during cellular senescence process (Hoare *et al.*, 2016; Teo *et al.*, 2019). In this study, we identified that cellular senescence caused an increase in Notch ligands expression in ECs. Given that the treatment with DNA methylation inhibitor, 5-AZA, abrogated the increased Notch ligands expression, senescence-associated epigenetic modifications might be critically involved in the dysregulated Notch ligands expression in senescent ECs. Further analyses for global DNA methylation status and histone modifications that affect the chromosome opening and closing are required to elucidate the mechanisms underlying the senescence-associated Notch signaling modulation.

One of the histopathological features of PAH-associated vascular remodeling is a remarkable medial thickening of distal pulmonary arteries that are normally non-muscularized. The pathological medial thickening is derived by pre-existing SMCs, which undergo dedifferentiation, distal migration, proliferation, and re-differentiation process (Sheikh, Lighthouse and Greif, 2014). These phenotypic plasticities of SMCs are tightly regulated by Notch3 signaling (Morrow *et al.*, 2008). Furthermore, Notch3 and its target gene HES5 are highly expressed in lung SMCs of PAH patients, and their expression levels are related to disease severity (Li *et al.*, 2009). Also, the Notch3-positive SMCs population has been identified as the origin of occlusive neointimal lesions of PAH (Steffes *et al.*, 2020). Of note, pharmacological and genetical inhibition of Notch3 signaling diminished medial thickening and ameliorated PH in mice (Li *et al.*, 2009; Steffes *et al.*, 2020). Therefore, Notch3 expressed in SMCs might be a major receptor for Notch ligands expressed in senescent ECs. However, further analysis is certainly needed to identify a critical Notch receptor(s) for senescence-associated Notch activation in SMCs through the interaction with ECs. Our study further emphasized the crucial role of Notch signaling in the pathogenesis of PAH, and newly revealed a crucial role of senescent ECs-SMCs interaction in the progression of pulmonary hypertension.

## Material and methods

### Animal study

All experimental protocols were approved by the Ethics Review Committee for Animal Experimentation of Kobe Pharmaceutical University. Transgenic mice that overexpressed TRF2DN in EC (VEcad-TRF2DN-Tg) were generated (C57/BL6J background). Mice were maintained under standard conditions with ad libitum to food (normal chow diet containing 23.1% protein and 5.1% fat) and water.

Mice at 8–9 weeks old were regularly used for experiments. For chronic hypoxia exposure, mice were put in the chamber with non-recirculating gas mixture of 10% O_2_ and 90% N_2_ for 3 weeks. For DAPT experiments, vehicle (10% ethanol and 90% corn oil) or DAPT (10 mg/kg dissolved in 10% ethanol and 90% corn oil) was injected subcutaneously, 3 times a week, during chronic hypoxia exposure, in accordance with previous publications (Cheng *et al.*, 2014; Sharma *et al.*, 2019).

### Echocardiography

Transthoracic echocardiography was performed using a Siemens Acuson X300 connected to a VF13-5SP probe (Siemens) to visualize the heart. The heart rate, left ventricular end-diastolic diameter, left ventricular end-systolic diameter, aortic diameter, pulmonary artery accelerated time (PAAT) and aortic velocity-time integral were measured. Three measurements were taken for each parameter and averaged. The ejection fraction and CO were calculated using the respective formula.

### Blood pressure measurements

The blood pressure was measured with a tail-cuff method using BP-98A-L (Softron) in a 37°C warmer without anesthesia. Five consecutive measurements were averaged. The results were presented in units of mmHg.

### Right ventricular pressure measurements

Right heart catheterization was performed with 1.4 F Millar mikro-tip catheter transducer under anesthesia using ~2% isoflurane inhalation. The catheter was inserted into right ventricle through right jugular vein. The right ventriclar systolic pressure was calculated from the average of five pressure waves and presented in units of mmHg.

### Fulton index measurements

The heart was dissected after 24–48 h fixation in 4% paraformaldehyde at 4°C. The right ventricle wall was separated from the left ventricle and septum and weighed separately. The data were presented as a ratio of the right ventricle to the left ventricle + septum.

### Histological analysis

The lung was inflated and fixed in 4% paraformaldehyde, followed by paraffin embedding. The sections were cut into 4 μm and stained with Elastica van gieson to quantify the distal pulmonary arteries (< 50 μm in diameter). The five randomly selected images of terminal bronchioles (20x magnification) was taken and the number of distal pulmonary arteries in 100 alveoli adjacent these terminal bronchioles were assessed. Five random images were captured using Keyence BZ-X800 fluorescence microscopy (Keyence).

Immunostaining was used to quantify the distal pulmonary artery muscularization. The lung sections were deparaffinized, followed by incubation in Antigen Unmasking Solution (Vector Laboratories) at 90°C for 10 min. The sections were blocked in 5% skim-milk in PBS with 0.2% Triton-X prior to incubation with anti-von Willebrand factor (Abcam) and FITC-labeled anti-α-smooth muscle actin (Sigma-Aldrich) antibodies at 4°C overnight. Subsequently, the sections were incubated with fluorescence-labeled donkey anti-rabbit secondary antibodies (Invitrogen), followed by mounting with Vectashield mounting medium with DAPI (Vector Laboratories). The 6 randomly selected images of distal pulmonary arteries (20x magnification) were taken using Keyence BZ-X800 fluorescence microscopy (Keyence) and used for quantification. The distal pulmonary arteries were determined as non-, partially-, and fully muscularized by positive α-smooth muscle actin staining < 25%, 25– 75%, and > 75% of the circumference, respectively. The data were presented as percent of non-, partially-, or fully muscularized PAs normalized with total number of vessels.

To assess the proliferation capacity of SMCs in distal pulmonary arteries, the immunostaining was performed as mentioned above using anti-Ki-67 (Histofine) and anti-α-smooth muscle actin (Sigma-Aldrich) antibodies. The data were presented as percent of fully muscularized distal pulmonary arteries with Ki-67-positive SMCs.

### Endothelial cell isolation

Isolation of mice lung endothelial cells was performed using gentle MACS Dissociator (Miltenyi Biotec Inc.), as indicated by the manufacturer. Briefly, 10–11 weeks old mice lungs were harvested, washed twice in PBS, cut into small pieces, and incubated in enzyme mix (Miltenyi Biotec Inc.). After 30 min incubation, homogenization of lungs were performed using gentleMACS Dissociators (#130-095-927, Miltenyi Biotec Inc.). The homogenate was filtered sequentially through 70 μm MACS Smart Strainer and 30 μm MACS Pre-Separation Filter. The dissociated cells were incubated with FcR Blocking reagent for mouse (#130-092-575, Miltenyi Biotec Inc.) for 10 min at 4°C, followed by incubation with CD146 (LSEC) Micro Beads (#130-092-007, Miltenyi Biotec Inc.) for 15 min at 4°C. Subsequently, cells were applied to LS column (#130-042-401, Miltenyi Biotec Inc.) in the magnetic field of MACS separator (#130-042-301, Miltenyi Biotec Inc.). After washing with PEB buffer for three times, isolated ECs were collected.

### TRF2DN plasmid construction and retroviruses production

The plasmid containing the TRF2-ΔB-ΔM (deletion mutant lacking the N-terminal basic domain and C-terminal Myb domain) was obtained from Addgene (plasmid #2431), as previously described (Barinda *et al.*, 2020). The TRF2DN/pMSCVneo and GFP/pMSCVneo construct were transfected into GP2-293 packaging cells using Lipofectamine 3000 (Thermo-Fisher) alongside with PVSVG viral envelope construct, followed by changing the medium in 24 h. After an additional 24 h incubation, fresh growth medium was given, and incubated for another 24 h. Subsequently, the culture medium containing retroviruses was collected and stored at −80°C after removal of cell debris by centrifugation.

### Quantitative real-time PCR

RNAs were extracted from the lungs or cells using RNAiso Plus (TAKARA), followed by purification using NucleoSpin RNA Clean-Up kit (Macherey-Nagel). The cDNA was synthesized from 0.5 and 1 μg of total RNA for cells and tissues, respectively, using PrimeScript RT Reagent Kit with gDNA eraser (TAKARA). Quantitative real-time PCR was performed using LightCycler96 (Roche Applied Science) with FastStart SYBR Green Master (Roche Applied Science). The mRNA expression levels of the target genes were normalized to 18S levels, and presented in arbitrary units. Nucleotide sequences of the primers are shown in Supplementary table 1.

### Retroviral transfection in PAECs

Frozen stocks of the viruses were thawed immediately before use. PAECs at ~70% confluence were infected with retroviruses using 1:1-2 mixture of retroviruses-containing medium and fresh growth medium. After 24 h incubation, fresh growth medium was given, followed by another 24 h incubation before use for experiments. The cells infected with retroviruses carrying GFP with more than 70% transfection efficacy were used as control cells.

### Cell culture

Human pulmonary arterial endothelial cells (PAECs) and smooth muscle cells (PASMCs) were purchased from PromoCell. PAECs and PASMCs were cultured in HuMedia-EG2 (Kurabo) and Smooth Muscle Cells Growth Medium 2 (PromoCell), respectively. All co-culture experiments were performed using 1 μm pore cell culture inserts (Falcon) coated with 1% gelatin (Wako). Indirect and direct co-culture was performed as previously described (Miyagawa *et al.*, 2019). Briefly, indirect co-culture was established by seeding GFP- or TRF2DN-transfected PAECs on the cell culture plate, and PASMCs were seeding on the cell culture insert. Direct co-culture was established by seeding GFP- or TRF2DN-transfected PAECs on the bottom of cell culture inserts for 2 h, and then PASMCs were seeding on the opposite side of the cell culture insert. In some experiments, 10 μM DAPT dissolved in dimethyl sulfoxide (DMSO) or vehicle (DMSO) were added after overnight incubation of direct co-culture condition. After 48 h incubation in indirect/direct co-culture, SMCs were assessed for proliferation, migration, and apoptosis capacities. Cells at passage 5 until 8 were used for all experiments.

In some experiments, PAECs were treated with 10 μM 5-azacytidine (5-AZA) dissolved in DMSO or vehicle (DMSO) for 72 h. Subsequently, cells were assessed for Notch ligands expression using Real-time qPCR.

### Cell proliferation assessment by immunofluorescence staining

PAECs and PASMCs were plated in 12-well plate and 12-well cell culture insert, both indirect and direct co-culture conditions with seeding density of 5×10^4^ cells/well and 1.5×10^4^ cells/insert, respectively. PASNCs on cell culture inserts were fixed by 4% paraformaldehyde, followed by blocking with 5% skim milk for 1 h. The cells were then incubated with Ki-67 antibody (Histofine) for overnight at 4°C. Subsequently, cells were incubated with fluorescence-labeled donkey anti-rabbit secondary antibodies (Invitrogen), followed by mounting with Vectashield mounting medium with DAPI (Vector Laboratories). The five randomly selected images (20x magnification) were taken using Keyence BZ-X800 fluorescence microscopy (Keyence) and used for quantification. The data were presented as a percentage of Ki-67-positive nuclei.

### Modified boyden chamber assay

PASMCs on cell culture inserts were trypsinized and seeded (20,000 cells per well) onto 8 μm pore inserts (Falcon). The number of migrated cells per well were assessed after 4 h, as previously described (Rinastiti *et al.*, 2020).

### Apoptosis assay

PAECs and PASMCs were plated in 6-well plate and 6-well cell culture insert, both indirect and direct co-culture conditions with seeding density of 10×10^4^ cells/well and 6×10^4^ cells/insert, respectively. To induce apoptosis, 500 nM hydrogen peroxide was added to PASMCs for 3 h. Cells were then trypsinized, followed by lyse in lysis buffer (10 mM Tris-HCl and 1% SDS), and boiled at 100°C for 5 min before centrifugation. Protein concentration was measured using the DC assay (Bio-Rad).

Cells lysates were run on SDS-PAGE, and transferred onto a nitrocellulose membrane. Subsequently, the membranes were incubated with antibodies for cleaved caspase 3 (Cell Signalling Technology), total caspase 3 (Cell Signaling Technology), or GAPDH (Cell Signalling Technology). The chemiluminescence signals were detected by BioRad ChamiDoc XRS system (BioRad).

### Statistical analysis

All data were presented as the mean ± SEM. Statistical analysis was performed using Graphpad Prism 9. The difference between groups was analysed using student *t*-test and two-way ANOVA with Tukey’s post hoc test, as indicated in the figure legend. *P*-value < 0.05 was considered as statistically significant. The number of experiments and animals per group were indicated in the figure legends.

## Acknowledgement

R.R was supported by Indonesian Endowment Fund for Education (LPDP) during her study in the division of Cardiovascular Medicine, Department of Internal Medicine, Graduate School of Medicine, Kobe University.

---

**Supplementary Figure 1.**
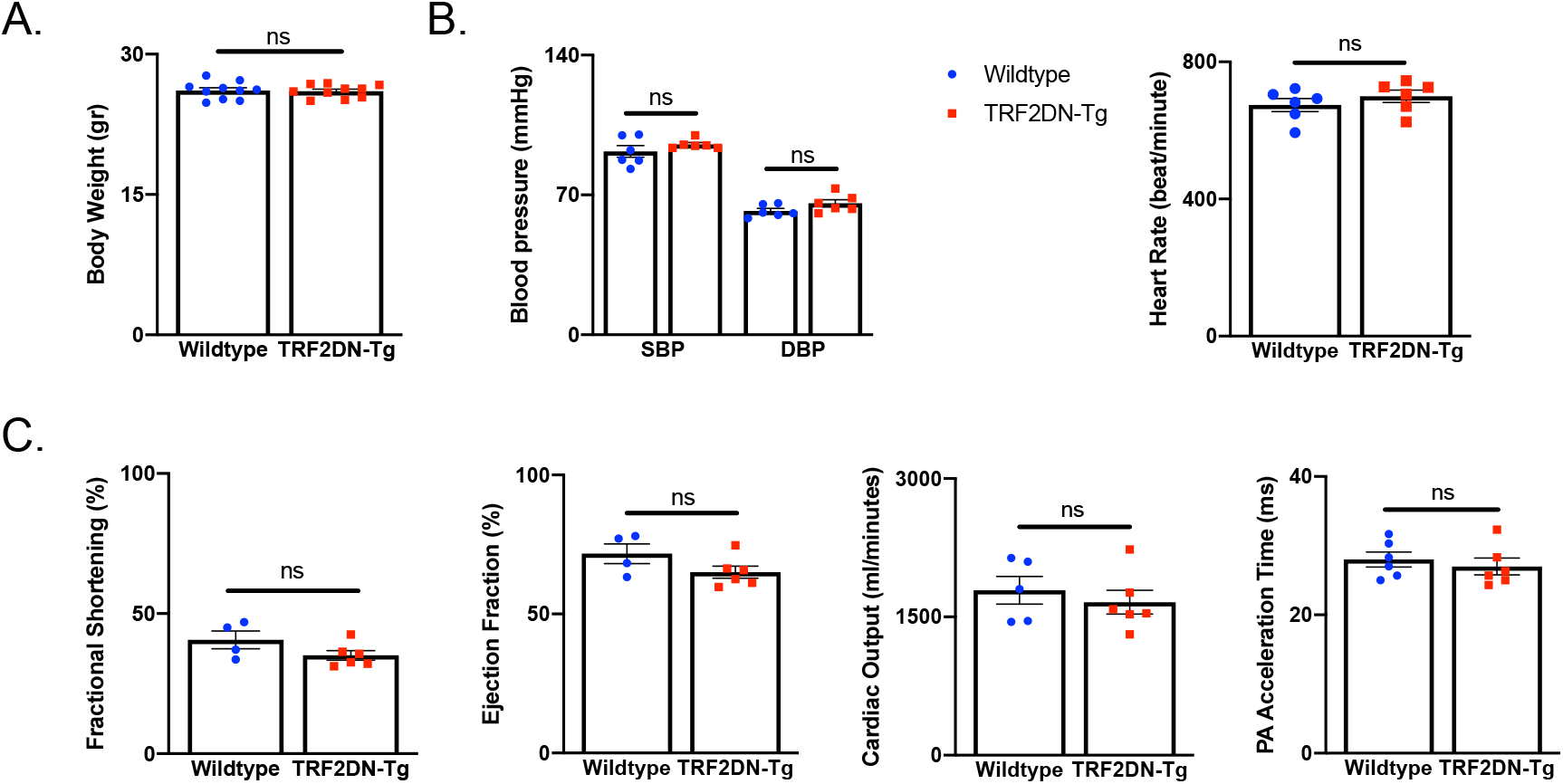
Basic characteristics of WT and EC-specific progeroid mice. (**A**) Body weight in WT and TRF2DN-Tg mice at the time of hypoxia exposure (n = 10 each). (**B**) Blood pressure and heart rate in WT and TRF2DN-Tg mice (n = 6 each). (**C**) Echocardiography parameters in WT and TRF2DN-Tg mice (n = 4-6 for WT; n = 5-6 for Tg). Data are presented as mean *±* SEM. Two-tailed student’s *t*-test was used for the analysis of the differences between two groups. *\*P < 0.05, **P < 0.01, ***P < 0.001, and ****P < 0.0001.*

**Supplementary Figure 2.**
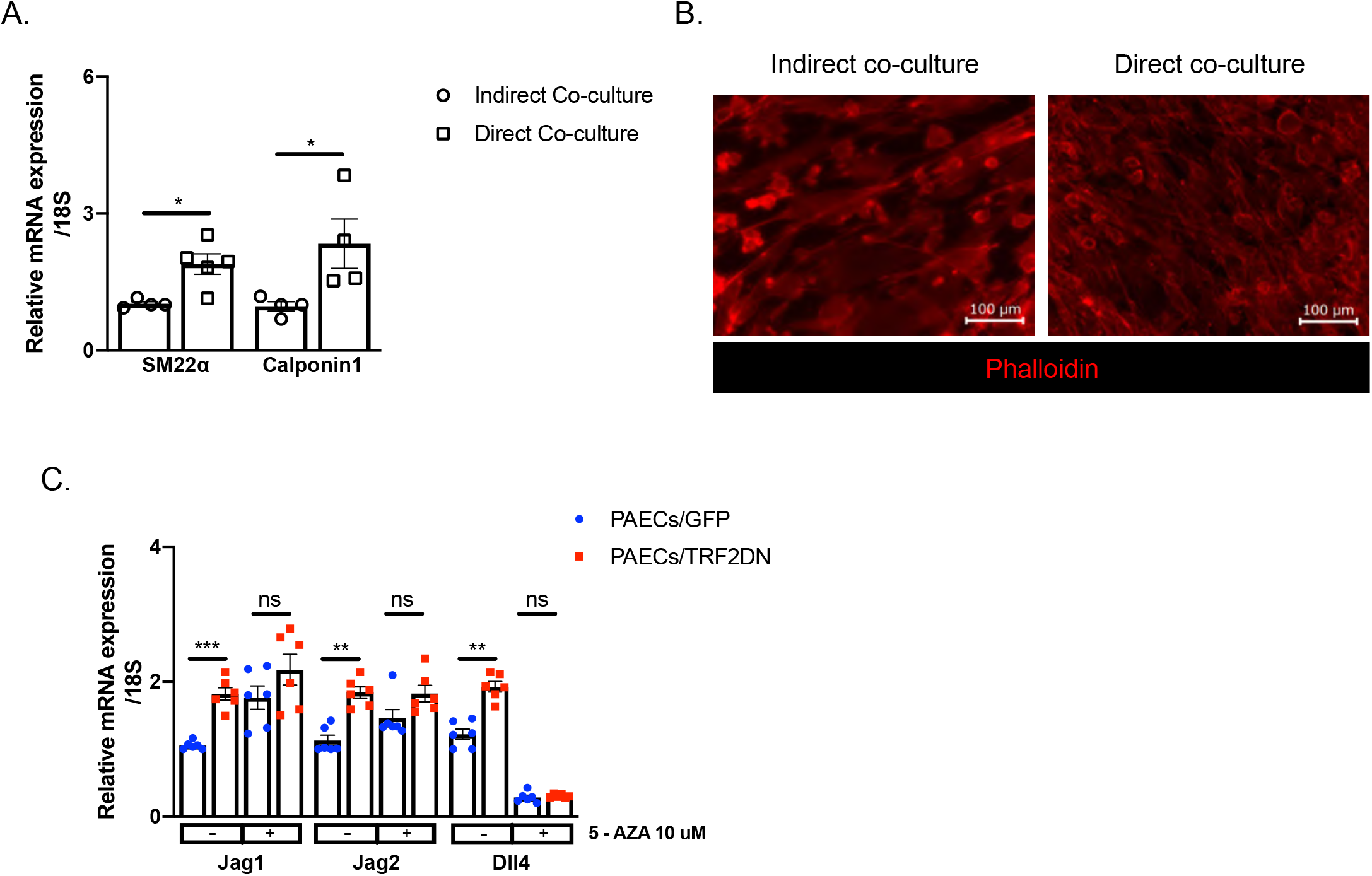
Characteristic of PASMCs and PAECs. (**A**) Real-time qPCR analysis for differentiation markers in PASMCs directly (n = 4 each) or indirect (n=4 each) co-cultured with PAECs. (**B**) Phalloidin staining in PASMCs directly or indirectly co-cultured with PAECs. (**C**) Real-time qPCR analysis for Notch ligands in PAECs transfected with either GFP (n = 6 each) or TRF2DN (n = 6 each). Cells were treated with with either vehicle or 10 μM 5-azacytidine. Data are presented as mean *±* SEM. Two-tailed student’s *t*-test was used for the analysis of the differences between two groups. Two-way ANOVA with Tukey’s post hoc test was used for the analysis of the differences between groups more than three. *\*P < 0.05, **P < 0.01, ***P < 0.001, and ****P < 0.0001.*

**Table S1.**
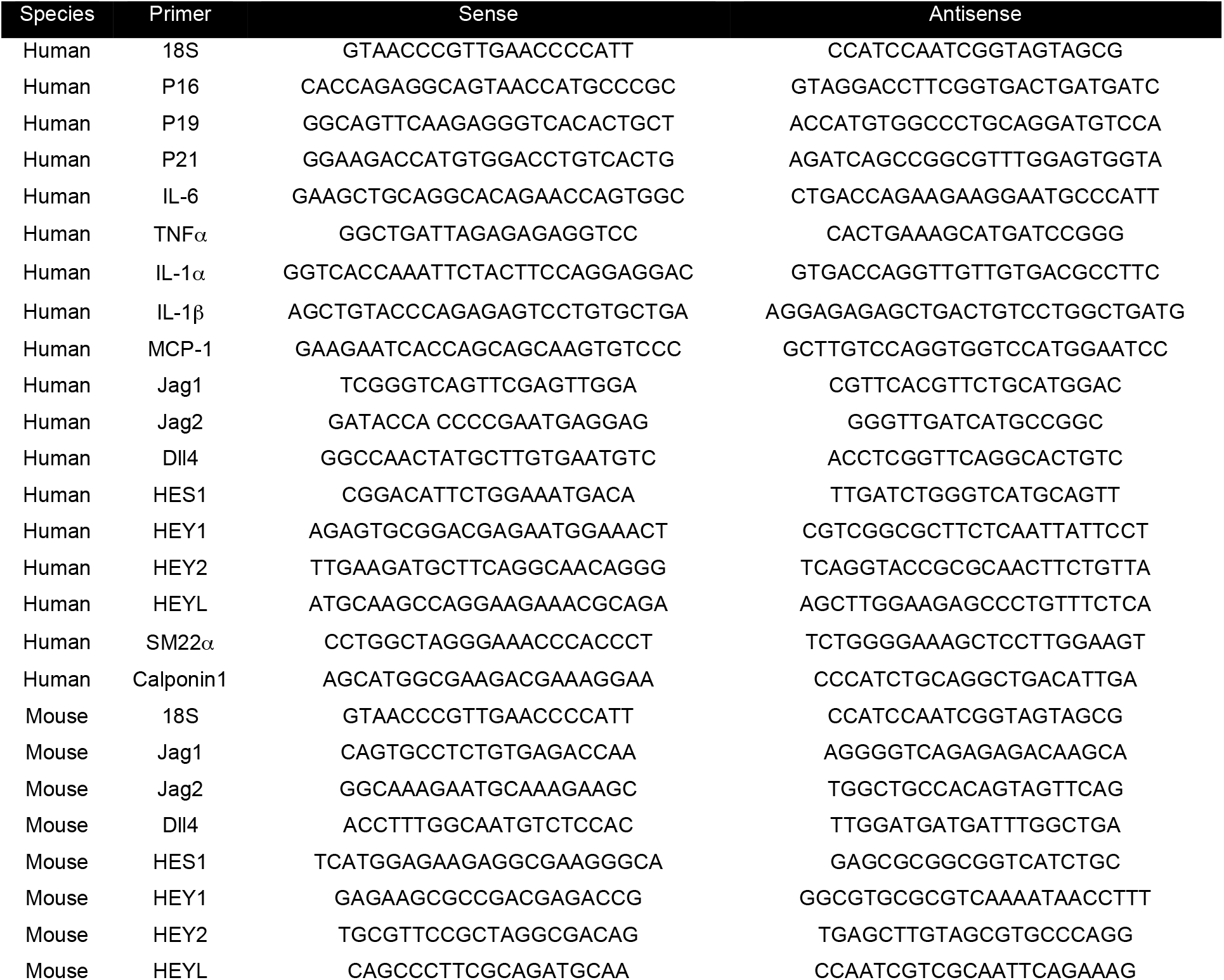
List of oligonucleotide sequences used for RTqPCR.

